# The Multi-State Epigenetic Pacemaker enables the identification of combinations of factors that influence DNA methylation

**DOI:** 10.1101/2023.01.24.525448

**Authors:** Colin Farrell, Kalsuda Lapborisuth, Sagi Snir, Matteo Pellegrini

## Abstract

Epigenetic clocks, DNA methylation based predictive models of chronological age, are often utilized to study aging associated biology. Despite their widespread use, these methods do not account for other factors that also contribute to the variability of DNA methylation data. For example, many CpG sites show strong sex-specific or cell type specific patterns that likely impact the predictions of epigenetic age. To overcome these limitations, we developed a multidimensional extension of the Epigenetic Pacemaker, the Multi-State Epigenetic Pacemaker (MSEPM). We show that the MSEPM is capable of accurately modeling multiple methylation associated factors simultaneously, while also providing site specific models that describe the per site relationship between methylation and these factors. We utilized the MSEPM with a large aggregate cohort of blood methylation data to construct models of the effects of age, sex and cell type heterogeneity on DNA methylation. We found that these models capture a large faction of the variability at thousands of DNA methylation sites. Moreover, we found modeled sites that are primarily affected by aging and no other factors. Among these, those that lose methylation over time are enriched for CTCF transcription factor chip peaks, while those that gain methylation over time are enriched for REST transcription factor chip peaks. Both transcription factors are associated with transcriptional maintenance and suggest a general dysregulation of transcription with age that is not impacted by sex or cell type heterogeneity. In conclusion, the MSEPM is capable of accurately modeling multiple methylation associated factors and the models produced can illuminate site specific combinations of factors that affect methylation dynamics.

## 1 Introduction

DNA methylation, the addition of a methyl group to the fifth carbon of the cytosine pyrimidine ring, is associated with the topological organization of the cellular genome, gene expression and the state of a cell. Within a population of cells the methylation pattern at certain sites can change predictably with the age of the individual from which the cells are drawn. This predictable nature of DNA methylation has led to the development of accurate DNA methylation based predictive models for age and health, termed epigenetic clocks. The difference between the predicted and the expected epigenetic age given an individual’s chronological age has been interpreted as a measure of age acceleration[1], and has been associated with mortality[2, 3] and other adverse health outcomes[4–8].

However, epigenetic clocks suffer from several limitations that limit the interpretability of their predictions and the underlying mechanisms. Epigenetic clocks are generally trained by using penalized regression based methods that attempt to minimize the difference between the predicted and observed value of age. As a result, as the error between predicted and observed age is decreased, the associations between age acceleration and mortality disappears[9]. Second generation epigenetic clocks attempt to resolve this issue by fitting a measure of human health, rather than age, and as a result these clocks are generally more sensitive to individual health status [10–12]. However, while the response variable is modified in these clocks the method used to fit the clock is largely the same. Epigenetic clocks are generally trained using regularized regression models, where the likelihood is maximized by minimizing the difference between the observed and predicted response variable subject to the elastic net penalty,*λ*_1_ and *λ*_2_. Methylation sites that increase model error and are influenced by other relevant factors such as smoking or obesity, may be discarded during model fitting, thus limiting the ability of this approach to account for the effects of these extraneous factors on epigenetic aging.

As an alternative to penalized regression based methods we previously developed an evolutionary based model for epigenetic dynamics, the Epigenetic Pacemaker (EPM)[13, 14]. The EPM attempts to minimize the difference between observed and predicted methylation values amongst a collection of sites through the implementation of a conditional expectation maximization algorithm[15]. Under the EPM the observed methylation status of a collection of sites is modeled linearly with respect to an input factor of interest, such as age. A hidden epigenetic state, that is related to the initial factor, but not necessarily linearly, is learned through the course of model fitting. The EPM can capture the non-linear relationship between methylation and age[16] and outputs an interpretable model for each site. However, both the EPM and regression based methods suffer from the same limitation, which is that they are limited to a single trait predicted by, or used to model, observed methylation patterns. In reality, the observed methylation landscape is likely impacted by a variety of factors that act simultaneously to produce the observed methylome of an individual.

To overcome this limitation, we have developed a multidimensional extension of the EPM, the Multi-State Epigenetic Pacemaker (MSEPM). We show that the (MSEPM) can accurately model site specific methylation variation driven by several factors, and given a trained model, accurately predict the values of the factors associated with an individual’s observed methylation profile in both simulated methylation datasets and a large aggregate blood tissue methylation dataset. Importantly, as factors that explain the observed methylation profile of an individual are added to the model the ability to model the factors and methylation values improves. Additionally, we show that sites with similar associations to modeled factors cluster together and are enriched for specific transcription factors.

## 2 Results

### 2.1 Multi-State Epigenetic Pacemaker Model

The MSEPM attempts to describe the observed methylation status at any single methylation site as a linear combination of methylation associated factors specific to an individual, termed epigenetic factors. Under this model epigenetic factors are related to observable individual factors, such as chronological age, sex and cell types, but may be transformed relative to observable factors. The epigenetic age factor, for example, often has a non-linear relationship with the observed age [16]. The MSEPM learns the appropriate transformation during model fitting to describe the observed methylation status linearly in terms of the epigenetic age factor (and not linearly with age). Given a site *i* and individual *j* the observed methylation status can be modeled as 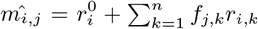, where *k* epigenetic factors, *f_j,k_*, are weighted by *k* site specific parameters, *r_i,k_*, and offset by a sites specific intercept term, 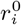. Site parameters, *r_i,k_* and 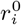, are characteristic of the site and shared amongst all individuals while epigenetic factors, *f_j,k_*, are characteristic of an individual and are the same across all sites for that individual. As a consequence, observed methylation differences between any two individuals are dependent on individual epigenetic factors *f_j,k_*. In practice, the observed methylation value is also dependent on a normally distributed error term 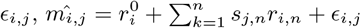.

Given an input matrix 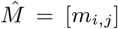 the objective of the MSEPM is to find the optimal values of *r_i,k_* and *f_k,j_* that minimize the residual sum of square (RSS) error, 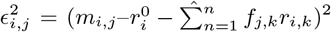. This is accomplished through the implementation of a conditional expectation maximization algorithm. The maximum likelihood (ML) values of *r_i,k_* and 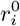 for each *i* methylation site can be solved using ordinary least squares (OLS) regression. Given the ML *r_i,k_* site coefficients each *k* epigenetic factor is then updated by fixing the site coefficients and updating *s_k,j_* by minimizing the RSS across all *i* sites using gradient descent, 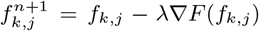, where *λ* is a specified learning rate. The optimization is accomplished by alternating between optimizing *r_k,i_* and *f_k,j_* until the reduction in sum of the site RSS is below a specified threshold between iterations or a set number of interactions is reached. Importantly, while the ML values of *f_j,k_* are by definition linear with the methylation status at any site, the original input factors for *f_j,k_* may not be.

Provided a trained MSEPM model and an unobserved methylation matrix, epigenetic factors are estimated by calculating each independent OLS for solution all *i* sites given the *r_i,k_* coefficients set for the respective input factor. These epigenetic factors can then be used to find the expected methylation value using the trained individual site models where 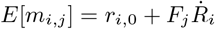.

### 2.2 Simulated Methylation Associated Factors

We implemented a simulation framework that extends the MSEPM model where the methylation status at site *i* for an individual *j* is described by a weighted sum of epigenetic factors, 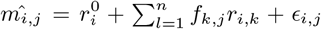. As shown in our previous work [16] the association between a methylation associated input factor and methylation status is not necessarily linear. To account for non-linear associations, *f_k,j_*, have the form 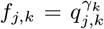, where *γ_k_* is the transformation between a factor of magnitude *q_n,j_* and the epigenetic factor. In practice the value of the *f_k,j_* is often unknown and the association between methylation status and *f_k,j_* is inferred through *q_n,j_*. We simulated individuals whose methylation is determined by four factors and their associated epigenetic factors: a uniform factor approximating age with a non-linear association with methylation status 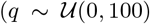, *s_Age_* = *q*^0.5^, Figure 1A-B), a binary trait resembling sex, linearly associated with methylation status (*q* ~ *B*(1, .5), *s_Sex_* = *q*, Figure 1C-D), a continuous normal (CN) phenotype resembling a cell type with a linear association with methylation status 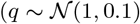, *s_CN_* = *q*, Figure 1E-F), and a continuous exponentially (CE) distributed trait resembling obesity with a linear association with methylation status (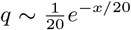, *s_CE_* = *q, FigureG* − *H*). Methylation sites were simulated by first randomly setting the dynamic range of the methylation site, (−1 *< δ* < 1), a site intercept, 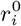, and the site error, 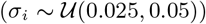. Simulated methylation sites were associated with a combination of zero, one, or multiple epigenetic factors. Rates for sites associated with multiple factors were set by sampling from a uniform distribution. Site rates were normalized to describe the dynamic range of the simulated methylation site unless the rates for all phenotypes were zero.

**Figure 1:**
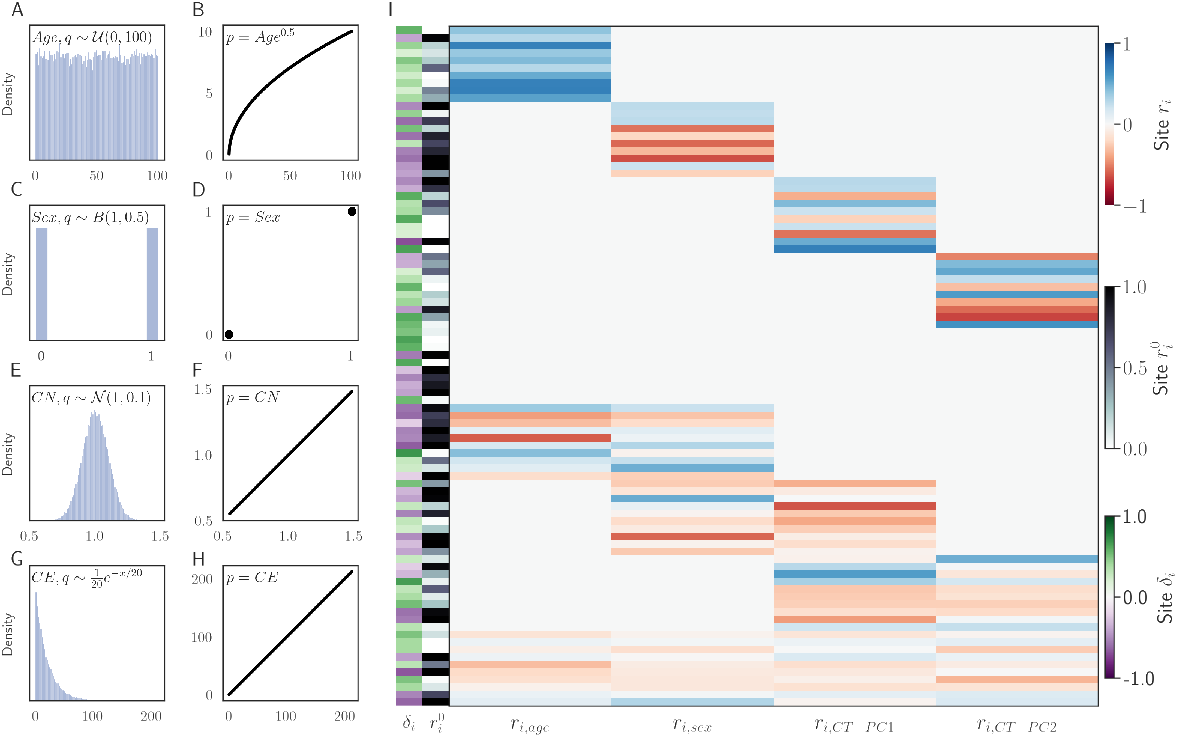
Simulated factors and the association with simulated methylation values. (A) Age with a non-linear association with methylation (B). Sex (C) with a binary association with methylation (D). Normal factor (E) with a linear relationship with methylation (F). Continuous exponential trait (G) with a linear relationship with methylation. (I) Simulated methylation sites. Each simulation site has a starting methylation value 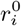, rate of change associated with each simulated factor *r_i,factor_* and range of variation *δ_i_*.

We simulated 90 methylation sites (Figure 1I). We then evaluated the MSEPM model as follows. We simulated 1000 samples with the four epigenetic factors described above. We then simulated methylation values using the simulated site rates. Simulated samples were then split for training (*n* = 500) and testing (*n* = 500). MSEPM models were then fitted using the values of the input factors, *q_k,j_*, as the initial guess for *f_k,j_*. We generated 1000 simulated datasets and fit MSEPM models using four combinations of input factors (Age, Age-Sex, Age-Sex-CN, Age-Sex-CN-CE). Within each simulation, epigenetic state predictions and methylation site predictions were made for all testing samples. All models captured the nonlinear association between simulated age and methylation (S. Figure 1). As the number of factors in the model is increased the mean absolute error (MAE) between the predicted epigenetic states and the simulated epigenetic factors decreases (Figure 2A). Importantly, to accurately assess simulated age it is necessary to account for the influence of the other simulated factors (Sex, CN, CE). The MAE between the predicted and simulated methylation values decreases as simulated factors are added to the model, and accurately assessing the methylation status of a simulated site requires that the factor associated with the methylation status at the site is included in the model (Figure 2A).

**Figure 2:**
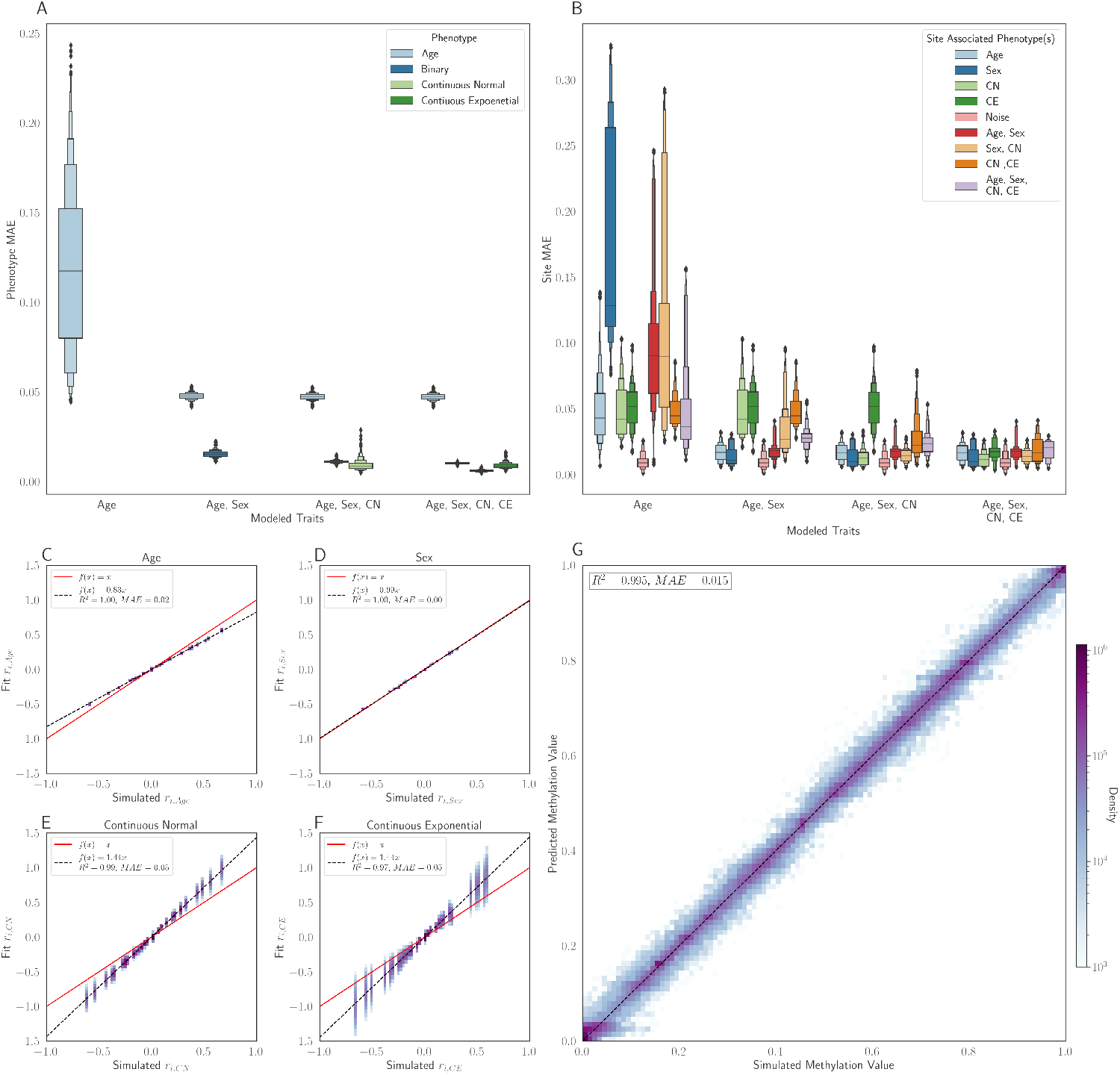
(A) The MAE of the factor predictions on the testing set as multiple factors are modeled simultaneously. (B) Prediction methylation MAE as factors are included in the MSEPM model. (C) Model coefficients for Age, Sex, Continuous Normal and Continuous Exponential factors for models trained (*n* = 500) with all four simulated factors. (D). Simulated and predicted methylation values for all simulated testing sites across all training folds.

The MSEPM model generated using all four simulated factors can capture the relative magnitude of the simulated site-specific rates (Figure 2C-F). However, the model has difficulty capturing the exact relationship between the simulated factors (age, CN and CE) and the inferred factors (Figure 2C, E-F). This is likely due to limitations of the model at capturing nonlinear methylation association and a limited training range for normally and exponentially distributed traits. Regardless, the four-factor model can accurately predict the simulated methylation value (Figure 2 D) and site intercept (Supp. Figure 1A). We also assessed the model robustness to variation in the number of samples and sites used for model training by randomly selecting a reduced subset of samples or sites for model training. MSEPM models trained with age, sex, CN, and CE can accurately assess all simulated phenotypes with few samples and sites (Supp Figure 1B-E).

### 2.3 Blood MSEPM Model

We utilized a large aggregated dataset composed of Illumina 450k array data from 17 publicly available datasets [7, 17–32] deposited in the Gene Expression Omnibus [33] (GEO) generated from blood derived samples (whole blood, peripheral blood lymphocytes, and peripheral blood mononuclear cells). All datasets were processed through a unified pipeline from raw array intensity data (IDAT) files using minfi (Aryee et al., 2014). Sex and blood cell type abundance predictions were made for each as previously described[34, 35]. The aggregate data spanned a wide age range (0.0 - 99.0 years, Figure 3A), contained more predicted females (*n* = 3392) than males (*n* = 2295, Figure 3B) and produced reasonable cell type abundance estimates (Figure 3C).

**Figure 3:**
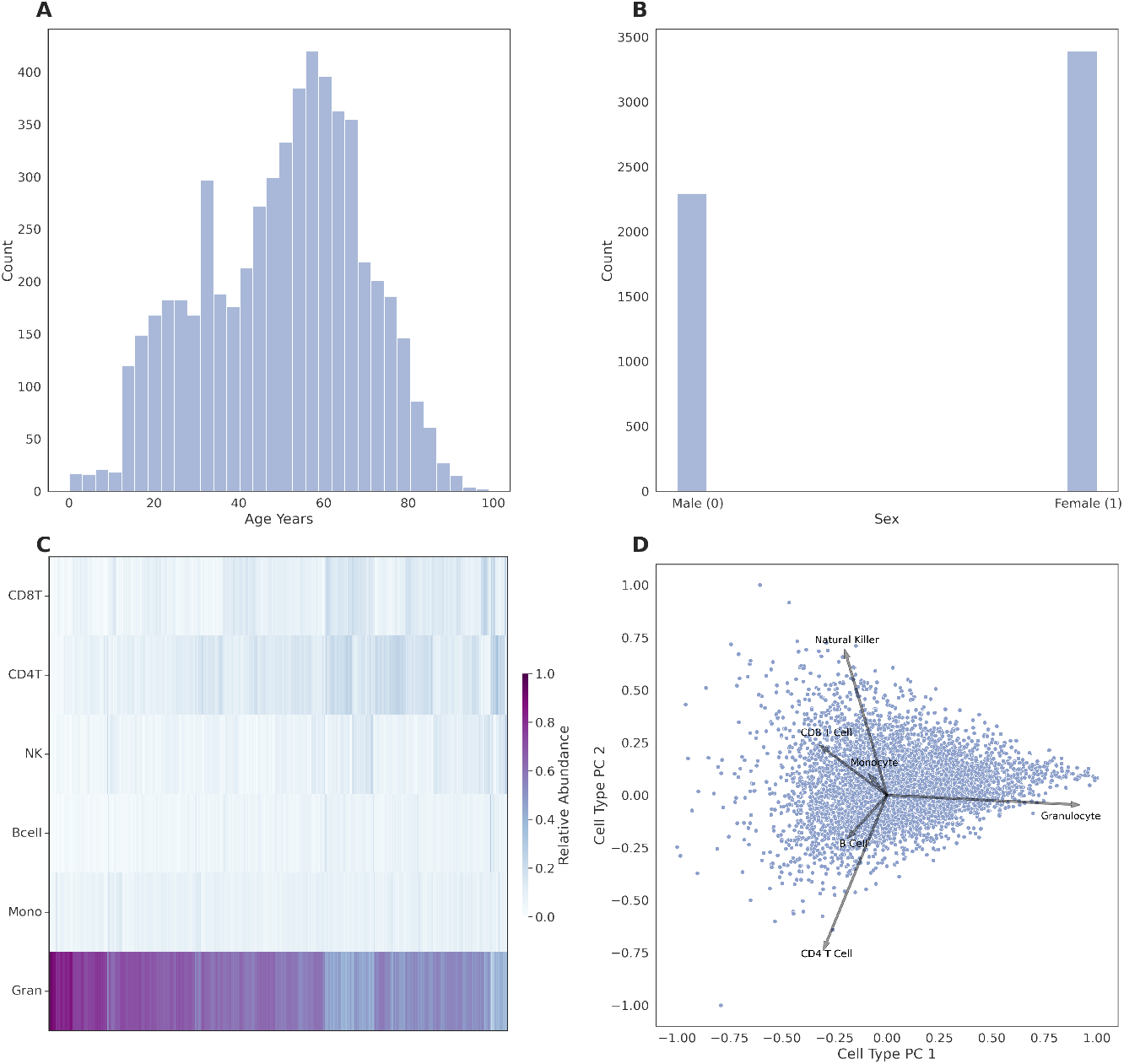
Distribution of age (A) and (B) sex in aggregate blood dataset. (C) Calculated cell type composition and (D) loading plot of principal components of cell type composition in aggregate blood data set. .

We trained MSEPM models using data assembled from four GEO series[20, 22, 29, 36] (*n* = 1605) with samples spanning a wide age range (0.01 - 94.0 years). The samples were randomly split into training (*n* = 1203) and validation (*n* = 402) sets stratified by age. Training set blood cell type abundance estimates were used to train a principal component analysis (PCA) model which was then used to calculate cell type PCA estimates for the validation and testing sets. The first cell type principal component (CT-PC1) is largely driven by the relative abundance of granulocytes (Figure 3D), while the second PC (CT-PC2) captures relative differences in the abundance of differentiated lymphocytes (Figure 3D). Methylation values for all samples were quantile normalized by probe type [37] using the median site methylation values across all training samples for each methylation site.

Methylation sites were selected for modeling with MSEPM if the sites were highly correlated with age (*n* = 276), sex (*n* = 49), CT-PC1 (*n* = 120), CT-PC2 (*n* = 116) or a combination of factors (*n* = 238), a total of 778 unique sites. Four MSEPM models were fit using the selected sites with four combinations of input factors (Age, Age Sex, Age Sex CT-PC1, and Age Sex CT-PC1 CT-PC2). Training sample factors were min-max scaled between 0 and 1 before model training. The association between the fit epigenetic factor predictions against the input modeled factors was assessed by fitting a trendline between epigenetic state predictions and scaled continuous input factors using the state prediction made for the MSEPM model trained with all four input factors. Performance of the MSEPM model was then evaluated using the testing samples (*n* = 4, 082). The performance of the MSEPM largely closely resembles the simulation results. All four MSEPM models capture the nonlinear relationship between age and methylation status (Supp. Figure 2). The epigenetic state prediction associated with age improves as the underlying methylation data are more fully explained through the addition of epigenetic factors (Supp. Figure 2). The MSEPM model fit with Age, Sex, CT-PC1 and CT-PC2 can accurately model the associated epigenetic state for each factor (Figure 4 A-D) and accurately predicts the methylation levels at individual sites (*R*^2^ = 0.935, *MAE* = 0.035, Figure 4 E). The trained MSEPM produces a collection of methylation site models that can help explain the association between modeled factors and methylation status.

**Figure 4:**
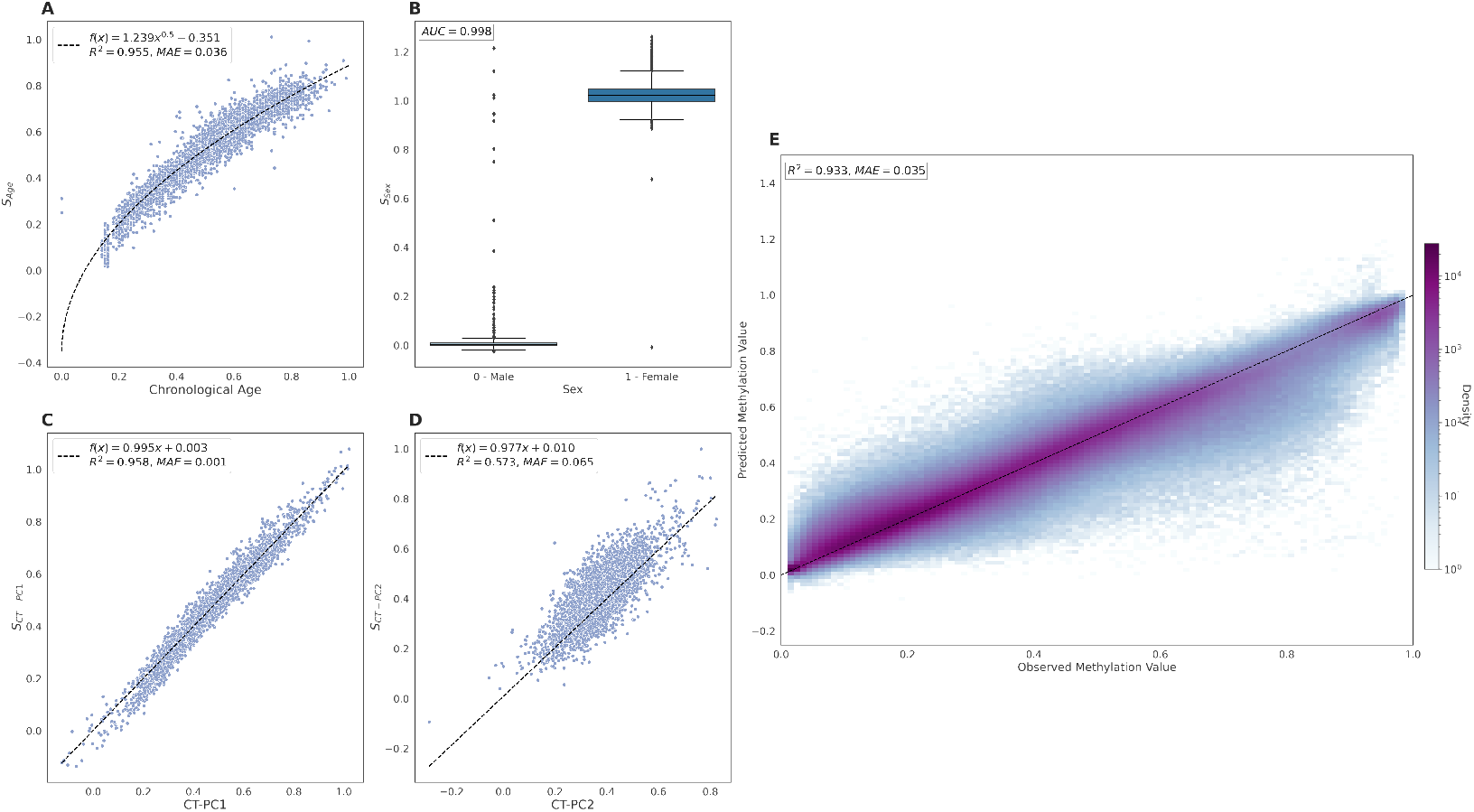
MSEPM model trained with age, sex, CT-PC1 and CT-PC2 predictions within testing set for methylation associated factors (A) age, (B) sex, (C) CT-PC1 and (D) CT-PC2. (E) Observed and predicted methylation values for training set has high concordance (*R*^2^ = 0.833, *MAE* = 0.035)

### 2.4 Analysis of chromatin regulators of site clusters

We evaluated the relationship between sites that are influenced by age, sex, CT-PC1 or CT-PC2 and potential regulatory factors by performing overlap enrichment analysis of these sites with transcription factor chromatin immunoprecipitation peaks present in the ENCODE V4 [38, 39] release. We first identified sites with similar coefficients of epigenetic factors through hierarchical clustering. Briefly, sites were clustered using Ward’s method by the euclidean distance between site regression coefficients that were normalized by the standard deviation of the methylation values amongst the training samples. The resulting tree was cut at a height of 18 to produce 10 distinct clusters with clear associations to the modeled factors (Figure 5A).

**Figure 5:**
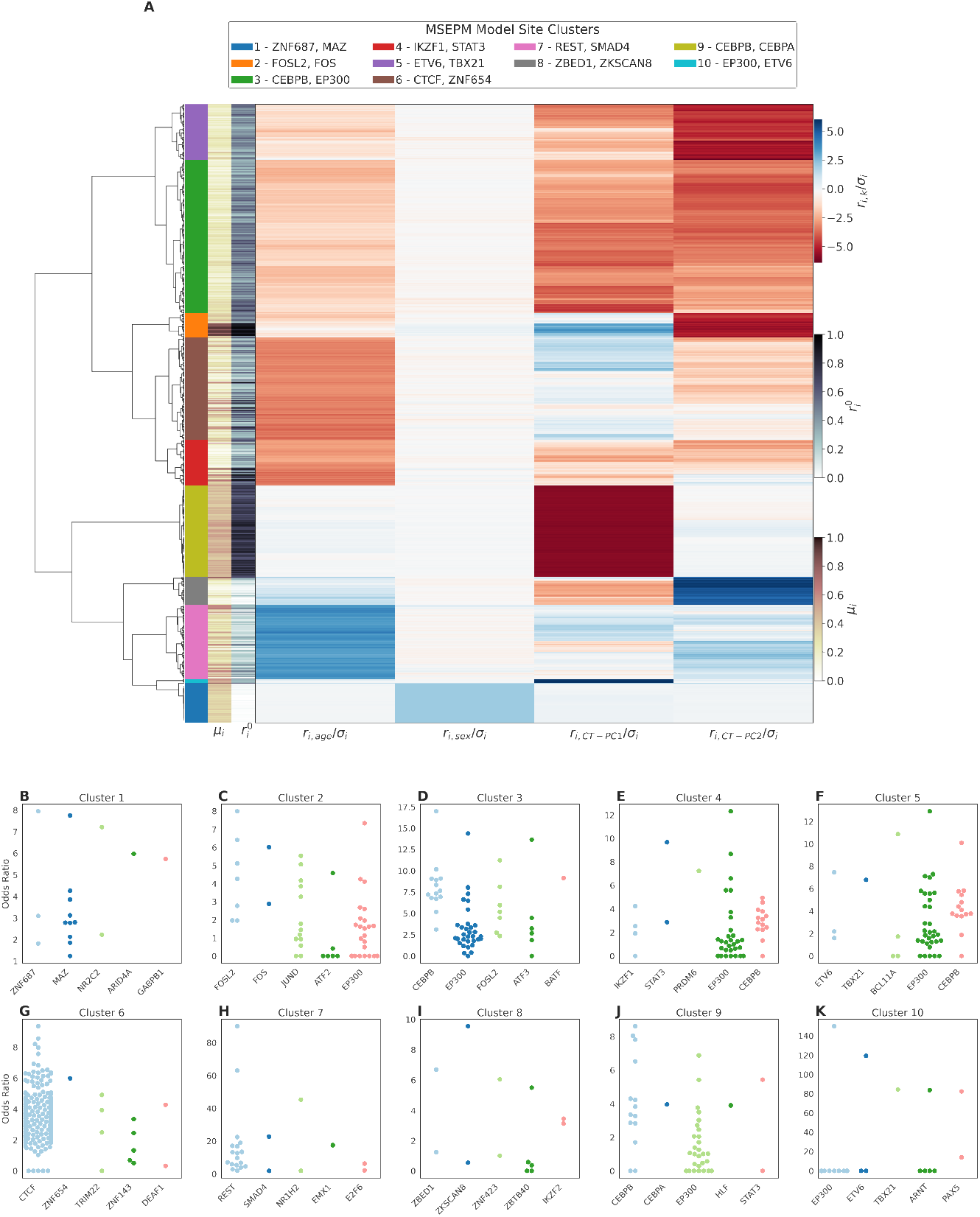
(A) Site clustering by standardized model coefficients. Sites clusters show distinct relationships with modeled traits. (B-K) Top five enriched transcription factors for clusters 1 - 10.

The site clusters largely conform to underlying biological expectations. Cluster one contains sites that are wholly associated with sex status and localized to the X chromosome (Supp. Table 1) and is enriched for peaks of transcription factors associated with sex specific regulation such as MAZ [40]. Clusters nine and ten contain sites whose methylation status is largely driven by CT-PC1, and are enriched for transcription factors associated with granulocyte development (CEBPB, CEBPA, EP300, ETV6)[41] [42]. Similarly, clusters two, five and eight are associated with CT-PC2 and are enriched for transcription factor peaks associated with immune development (ZBED1, ETV6, FOSL2, FOS, TBX21). Clusters four and six are associated with loss of methylation with age. Cluster six is highly enriched for CTCF binding sites; CTCF is known to increase at sites where methylation is lost during aging [43]. Cluster four is enriched for STAT3 whose activation during exercise is age dependent [44, 45]. Cluster seven is associated with the accumulation of methylation with age and is enriched for immunoprecipitation peaks for aging associated transcription factors SMAD4 and RE1-Silencing Transcription Factor (REST). SMAD4 encodes a protein involved in the transforming growth factor beta (TGF-*β*) signaling pathway. Age related dysregulation of TGF-*β* has been linked to reduced skeletal muscle regeneration[46, 47] and SMAD4 polymorphisms are associated with longevity [48]. REST is a transcriptional repressor of neuron specific genes in non-neuronal cells [49, 50]. REST expression is upregulated in aged prefrontal cortex tissue and the absence of REST expression is associated with cognitive impairment [51] and cellular senescence in neurons [52].

## 3 Discussion

Epigenetic clocks are widely used tools to study human aging and health. Despite their widespread use, the biological interpretation of model parameters is limited. A methylome is influenced by many different biological processes occurring simultaneously over time that may differ among individuals. Epigenetic clocks, while producing accurate predictions of age, attempt to capture this complexity through a single dependent variable. Additionally, the penalized regression based methods used to fit most epigenetic clocks select sites that minimize, or regress out, the influence of other factors and omit groups of sites that are correlated. To overcome these limitations, here we propose a multidimensional extension of the EPM model, the MSEPM.

In contrast to previous methods, the MSEPM aims to simultaneously model the effect of multiple factors on the methylome.The simulation and blood MSEPM models show that concurrently modeling age, cell type composition and sex can minimize model residuals when compared with the MSEPM model fit with age only. The residual of the age only model is often interpreted as a measure of age acceleration. When multiple methylome associated traits are modeled simultaneously this residual can be explained directly by other factors and the association between the methylome and a trait of interest can be inferred.

Additionally, the individual methylation site linear models fit as part of the MSEPM optimization can provide information about the relationship between modeled factors and site specific biology. To this end, we find that the blood MSEPM model conforms to expected biology. Sites with a strong sex association localize to the X chromosome and sites associated with cell types are enriched for transcription factors associated with the development of immune cells. Sites that are primarily affected only by age in the blood MSEPM model are of particular interest. As others have previously described, sites that progressively lose methylation over time are strongly enriched for CTCF [53, 54], while in the blood MSEPM model sites that gain methylation over time are enriched for REST. The association between age and REST is well established, but the association between gain of methylation and age has previously not been reported. Both transcription factors are associated with the maintenance of transcriptional profiles and suggest that there is a general dysregulation of maintenance of cell specific chromosomal organization over time.

In conclusion, we introduced a multi-dimensional extension of the Epigenetic Pace-maker, the MSEPM. The MSEPM is capable of accurately modeling multiple methylation associated factors simultaneously. This paradigm can elucidate the site specific regulation underpinning methylome dynamics. The MSEPM is available under the MIT license at https://github.com/NuttyLogic/MultistateEpigeneticPacemaker.

## 4 Methods

### 4.1 MSEPM

MSEPM model training begins with an input matrix of *i* methylation sites for *j* individuals and *k* input traits for *j* individuals. The goal of the MSEPM is to minimize the difference between the observed and predicted methylation values at any of the *i* input methylation sites as explained by the *k* input traits. The MSEPM model is fit as follows:

- Fit *i* ordinary least squares regression models with the form 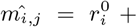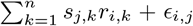 to estimate site specific parameters *ri* shared amongst all individuals.
  – The *k* untransformed input traits are used as the initial guess of *S_j,k_* for the first model iteration.
- Update *S_j,k_* to minimize the RSS cost function, 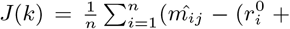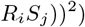, using gradient descent with fixed *ri* site parameters.
  – ∇*J* (*k*)

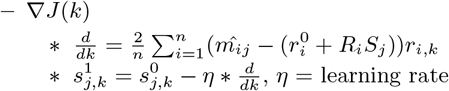
- Repeat steps 1-2 until the reduction in sum of site RSS between iterations is ≤ specified threshold or a set number of iterations has been reached.

### 4.2 Simulation Framework

Under the MSEPM formulation a single site can be described linearly where the observed methylation value is dependent on a weighted sum of *n* epigenetic states, 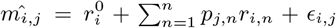, where *p_j,n_* is an individual trait associated with the methylation status at site *i*. Using this formulation we implemented a simulation framework in python with only numpy [55] as a dependency. The details of the approach are as follows:

- Individuals are simulated by setting individual traits with the following form 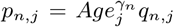 Where the simulated individual trait *p_n,j_* is dependent time and exposure, *qn,j*, over time. Under this formulation traits may have a non-linear association with the simulated methylation status as mediated by time. The form of *Q*[*q*] varies between simulated traits as shown above. Simulated MSEPM models are fit using *qn,j* as input for and the association between *pn,j* is learned through model fitting. 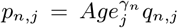 Where the simulated individual trait *pn,j* is dependent time and exposure, *qn,j*, over time. Under this formulation traits may have a nonlinear association with the simulated methylation status as mediated by time. The form of *Q*[*q*] varies between simulated traits as shown above. Simulated MSEPM models are fit using *qn,j* as input for and the association between *pn,j* is learned through model fitting.
- Methylation site profiles are then simulated by setting the dynamic range of possible methylation values and the error profile of the methylation site. The simulated methylation values are bounded by 0 and 1, and represent the proportion of methylated bases to the total number of observed cytosines at any site. The dynamic range is set by first initial methylation value 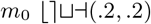 and then setting a target methylation value *m_t_* where the dynamic range is *δ_i_* = *m_t_* − *m_o_*. The value of *m_t_* is set conditionally to ensure dynamic range is always larger than some specified threshold, *θ*, where *θ* ≤ |*δ*| ≥.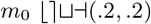 and then setting a target methylation value *m_t_* where the dynamic range is *δ_i_* = *m_t_* − *m_o_*. The value of *m_t_* is set conditionally to ensure dynamic range is always larger than some specified threshold, *θ*, where *θ* ≤ |*δ*| ≥.
- Methylation sites profiles are then randomly selected and associated with a weighted set of simulated traits. The weighted traits are then normalized so the input combination of traits describes the dynamic range of the simulation site, *δ*. If no weights are provided the individual methylation sites are simulated by returning *m*_0_ +*N* (*μ, σ*). *δ*. If no weights are provided the individual methylation sites are simulated by returning *m*_0_ + *N* (*μ, σ*).

### 4.3 Blood MSEPM Model Training

Methylation sites with an absolute pearson correlation coefficient greater than 0.7, 0.995, 0.92 and 0.64 between age (*n* = 276), sex (*n* = 49), cell type PC1 (*n* = 120) and cell type PC2 (*n* = 238) and site methylation values respectively among training samples samples(*n* = 1203) were included in the model. Additionally, sites with a sum of all absolute pearson coefficients greater than 1.8 were also included (*n* = 238) for a total of 778 methylation sites. Min-max, (0-1), scalers were fit using the training input features. Validation and testing sample features were transformed with the trained scalers, note age was min-max scaled on a range from 0-100 years. MSEPM models were trained with a learning rate of 0.01 with an iteration limit of 200.

### 4.4 Blood MSEPM Model Cluster Transcription Factor Overlap Analysis

A custom transcription factor reference set was built from all ENCODE V4 transcription factor chromatin immunoprecipitation [38, 39] (release 1.4.0 - 2.1.2) irreproducible discovery rate narrow bed peaks, which contains peaks with high rank consistency between replicates, that were not audited for non-compliance or errors. GRCh38 region coordinates were lifted to GRCh37 coordinates using liftOver [56]. The overlap reference contains 714 transcription factor targets 1621 accession IDs (table. . .).

Blood MSEPM model site hierarchical clustering was performed as follows. Individual methylation site coefficients were first normalized by the standard deviation of methylation values of the site among the training samples, *r_i,n_/σ_i_*. A distance matrix was then created by taking the Euclidean distance between the normalized site model coefficients. Sites were then clustered using Ward’s method which seeks to minimize within cluster variance by minimizing the increase in the error sum of squares (ESS) through successive cluster fusions. Clusters label by tree cutting at a height of 18. All clustering analysis was carried out using SciPy v1.6.3 [57]. *r_i,n_/σ_i_*. A distance matrix was then created by taking the Euclidean distance between the normalized site model coefficients. Sites were then clustered using Ward’s method which seeks to minimize within cluster variance by minimizing the increase in the error sum of squares (ESS) through successive cluster fusions. Clusters label by tree cutting at a height of 18. All clustering analysis was carried out using SciPy v1.6.3 (Virtanen et al. 2020).

Transcription factor enrichment analysis was performed with LOLA [58] which assesses the genomic region set overlap between a set of query regions and a set of reference regions, within a specified shared background set, using Fisher’s exact test. Overlap analysis was performed for sites within a cluster against the ENCODE V4 reference region (1BP minimum overlap) using all sites assayed with Infinium Human-Methylation450K BeadChip as background.

### 4.5 Analysis Environment

Analysis was carried out in a Jupyter[59] analysis environment. Joblib[60], SciPy[61], Matplotlib[62], Seaborn[63], Pandas[64] and TQDM[65] packages were utilized during analysis.

### 4.6 Supplementary Information

All analysis code, data processing code, and supplementary material associated with this manuscript can be found at https://github.com/NuttyLogic/MSEPMManuscript. The methylation simulation utility can be found at https://github.com/NuttyLogic/MethSim. The data supporting these findings are openly available at GEO under the series GSE87640, GSE87648, GSE51057, GSE51032, GSE87571, GSE125105, GSE42861, GSE69138, GSE111629, GSE128235, GSE121633, GSE73103, GSE61496, GSE59065, GSE97362, GSE156994, GSE128064 and GSE43976.

