## Supplementary Information for "The Multi-State Epigenetic Pacemaker enables the identification of combinations of factors that influence DNA methylation"

---

### 1 Supplementary Table Descriptions

**Supplemental Table 1:** MSEPM model parameters for MSEPM blood model trained against age, sex, CT-PC1 and CT-PC2.
**Supplemental Table 2:** Sample characteristics for GEO samples used in MSEPM blood model training, validation, and testing.
**Supplemental Table 3:** LOLA transcription factor binding results.
**Supplemental Table 4:** Simulated methylation site parameters.

### 2 Supplementary Figures

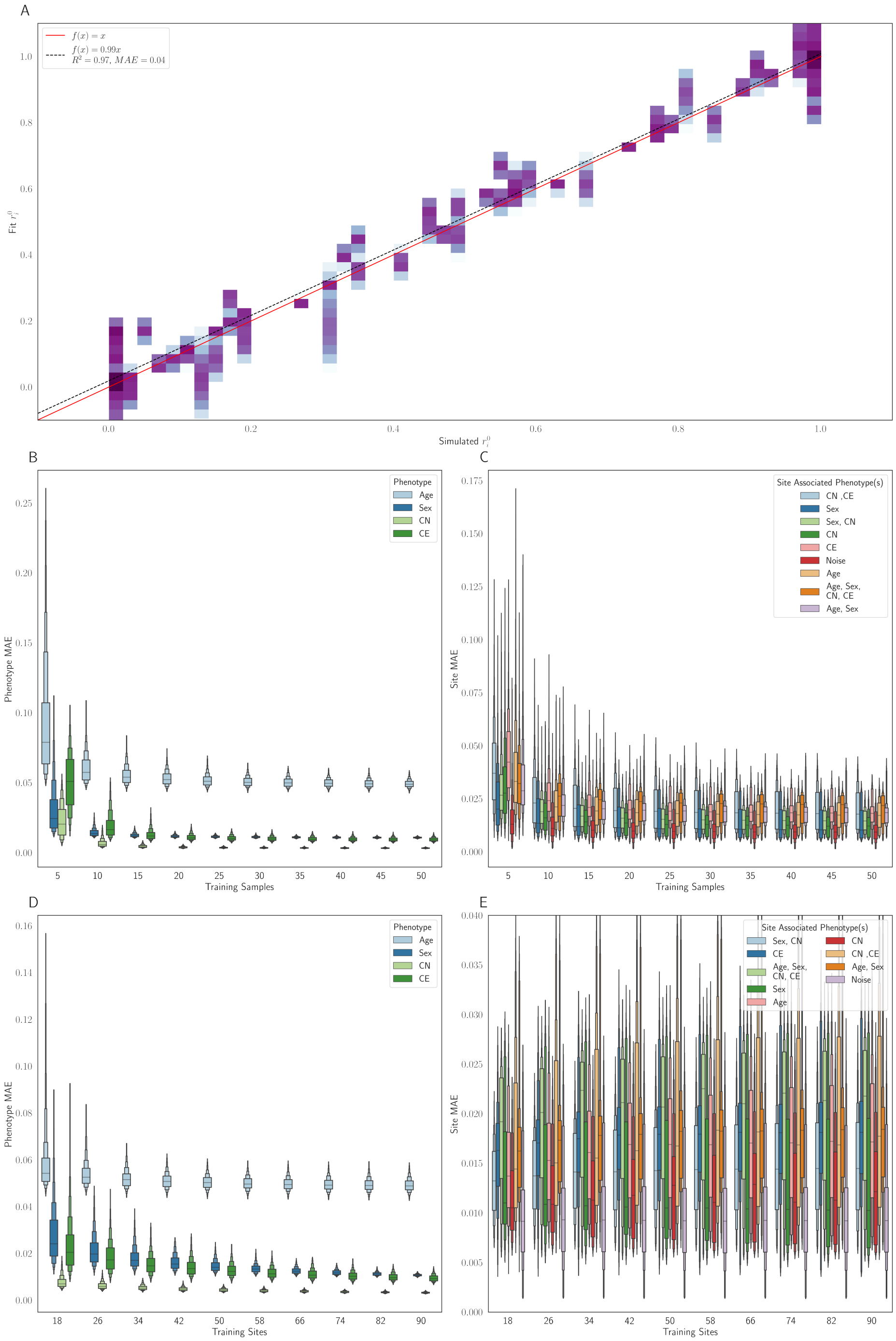


**Supp. Figure 1:** Simulated methylation site intercept accurately modeled with MSEPM four factor model (A). Simulation phenotype (B) and site methylation (C) prediction MAE for four-factor MSEPM models fit with a varying number of training samples and sites (D-E).

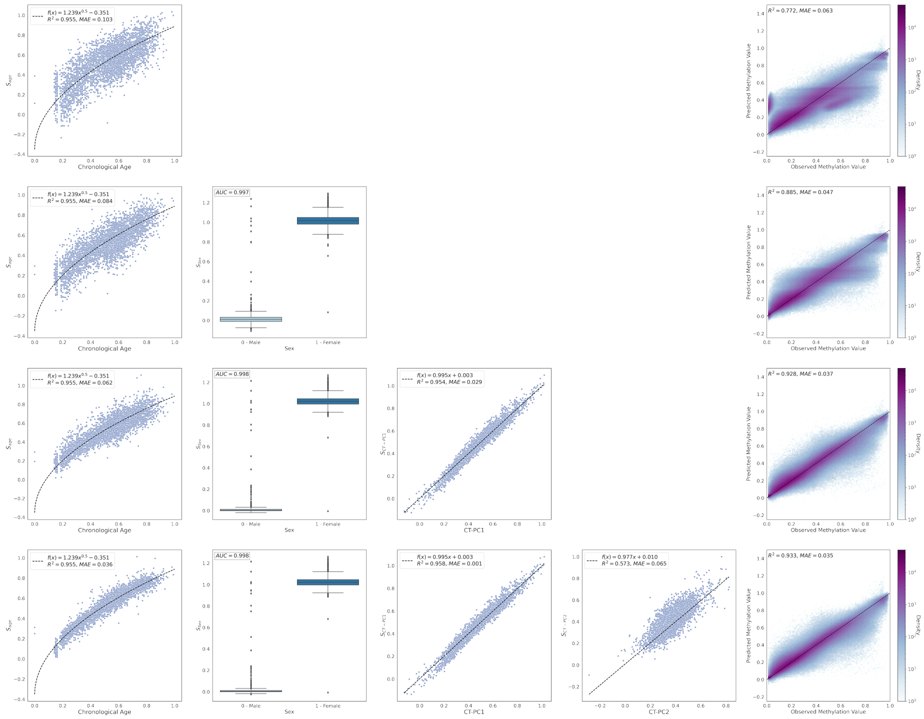


**Supp. Figure 2:** MSEPM testing blood model predictions for MSEPM model fit with only age (first row), age / sex (second row), age / sex / cell type PC1 (third row), and age / sex / cell type PC1 / cell type PC2 (fourth row).

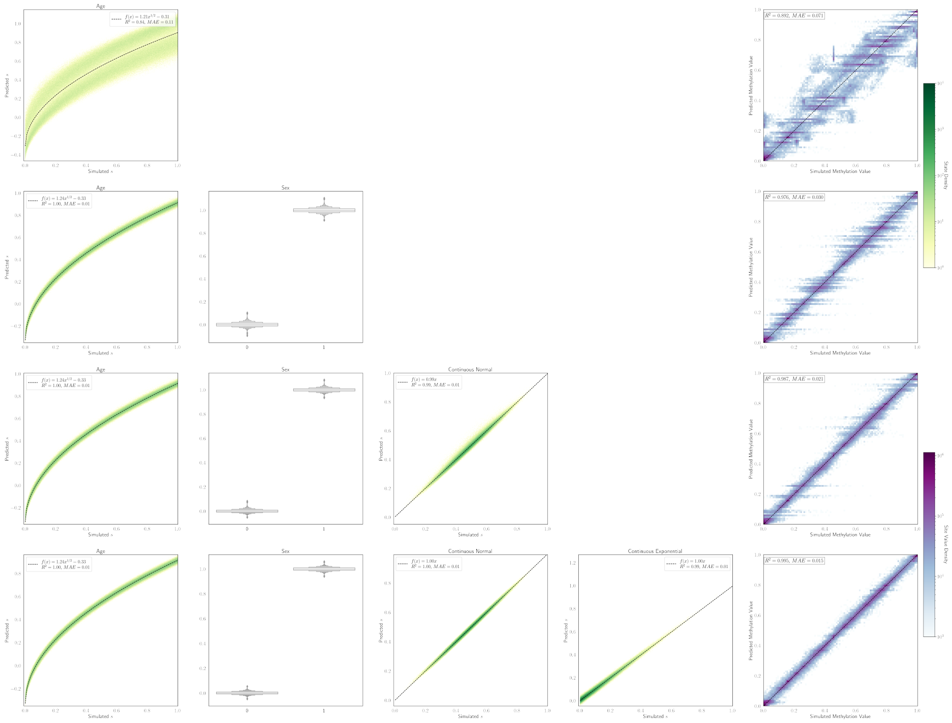


**Supp. Figure 3:** Simulated testing model predictions for MSEPM model fit with only age (first row), age / sex (second row), age / sex / CN (third row), and age / sex / cell type PC1 / cell type PC2 (fourth row).

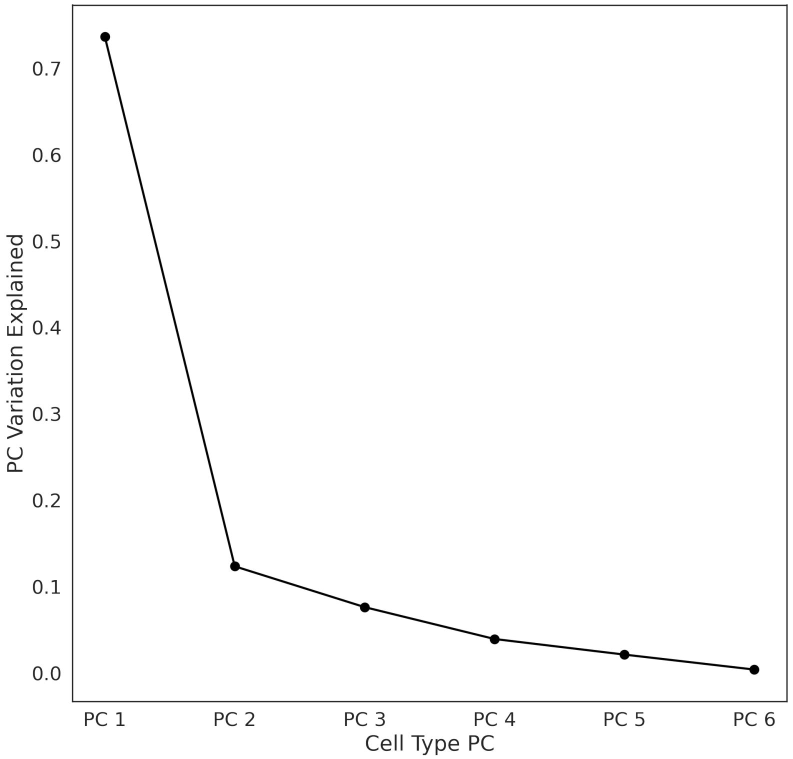


**Supp. Figure 4:** Cell type principal component analysis scree plot.

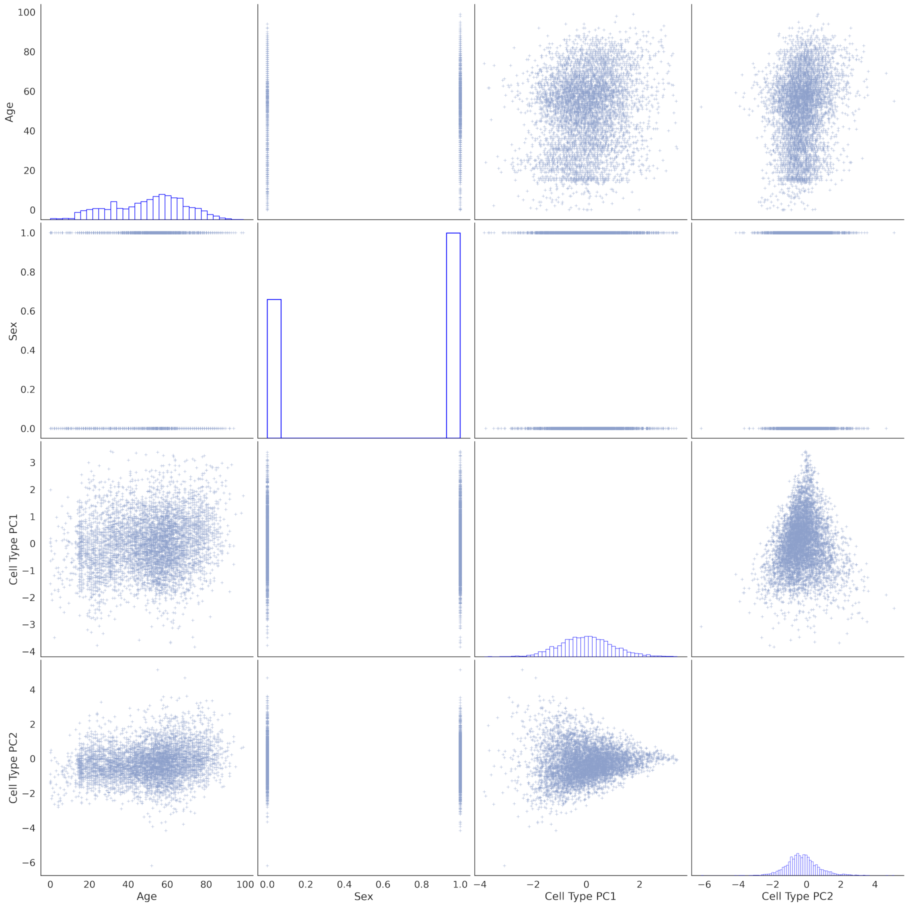


**Supp. Figure 5:** Pairwise bivariate distributions, and single factor distributions plots, for GEO factor data (age, sex, CT PC1 and CT PC2) used in MSEPM model training.

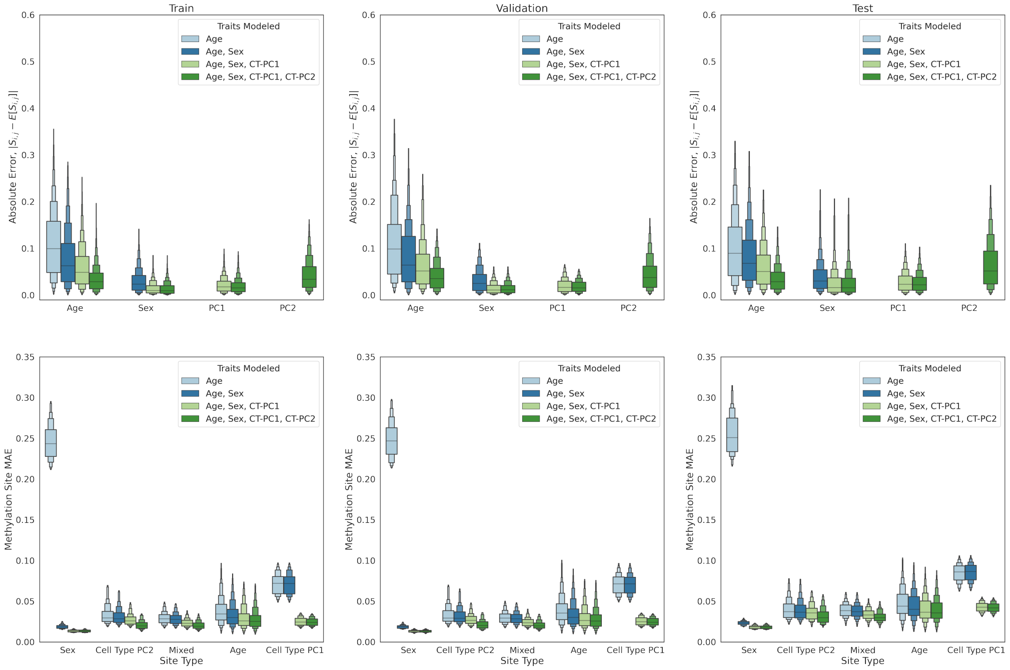


**Supp. Figure 6:** Methylation site and sample factor prediction error for models trained with 1 to 4 factors for training, validation and testing sets.

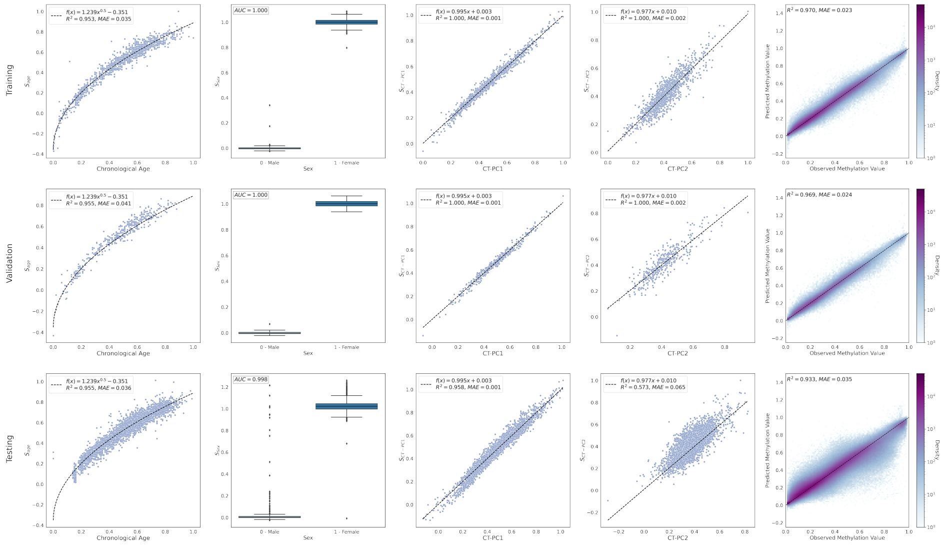


**Supp. Figure 7:** MSEPM blood model predictions for MSEPM model fit against age, sex, CT PC1 and CT PC2 for training, validation and testing sets.
